# Near-chromosome-level genome assembly and transcriptome analysis of the Ural owl, *Strix uralensis* PALLAS, 1771

**DOI:** 10.1101/2025.03.26.645461

**Authors:** Sven Winter, René Meißner, Martin Grethlein, Gerrit Wehrenberg, Angelika Kiebler, Andrea X. Silva, Natalia Reyes Escobar, Suany M. Quesada Calderón, Ana V. Suescún, Luis Guzman Belmar, Stefan Prost

## Abstract

The Ural owl (*Strix uralensis*) is a large member of the Strigidae family and inhabits Eurasian forests ranging from Germany to Japan. However, it faces increased range reduction, particularly at its southwestern distribution edges. Despite being considered “Least Concern” by the IUCN, local populations have become threatened in Central Europe due to severe habitat loss. Reintroduction programs aim to restore these populations by closing distribution gaps and facilitating natural recolonization of suitable habitats. To support these efforts, genomic resources have become an established tool to assess genetic diversity, geographic structure, and potential inbreeding, crucial for maintaining the genetic health and adaptability of newly established populations. Here, we present a *de novo* genome assembly and transcriptome of the Ural owl based on ONT long-reads, Omni-C Illumina short-reads, and RNASeq data. The final assembly has a total length of 1.26 Gb, of which 96.37 % are anchored into the 41 largest scaffolds. The contig and scaffold N50 values of 88.6 Mb and 21.7 Mb, respectively, a BUSCO/compleasm completeness of 97.5 %/99.65 % and k-mer completeness of 95.18 %, emphasize the high quality of this assembly. Furthermore, annotation of the assembly identified 17,650 genes and a repeat content of 12.48 %. This new highly contiguous and chromosome-scale assembly will greatly benefit Ural owl conservation management by informing reintroduction programs about the species’ genetic health and contributing a valuable resource to study genetic function in greater detail across the whole Strigidae family.

## Introduction

Owls (Strigiformes) are primarily nocturnal birds of prey, known for their exceptional low-light vision and excellent hearing capabilities. Unlike most orders of birds, owls primarily rely on their hearing for hunting, and acoustically detected prey is captured in an almost silent approach, made possible by specialized wing anatomy (Konishi, 1973). Today, the Strigiformes consist of approximately 250 extant species, classified into two distinct families: Tytonidae, or barn owls, with 19 species, and Strigidae, or true owls, with approximately 235 species (Ackerman, 2024; Penhallurick, 2002).

Within the Strigidae, the Ural owl *Strix uralensis* (Fig. 1) is a large member of the wood owls (*Strix*) with a round body and an exceptionally long tail (Voous, 1964). The species prefers old-growth primary forests and requires sufficient open areas without underbrush, allowing for unobstructed hunting, especially during the rearing of its young (Tutiš et al., 2009). Its usual prey ranges from small rodents to medium-sized lagomorphs and varies depending on the populated habitat, with the most frequently occurring prey being the most commonly hunted (Korpimäki & Sulkava, 1987). Throughout Eurasia, Ural owls exhibit a wide distribution across various forest habitats, extending from Scandinavia and parts of Central Europe to Japan (BirdLife International, 2021).

**Figure 1.**
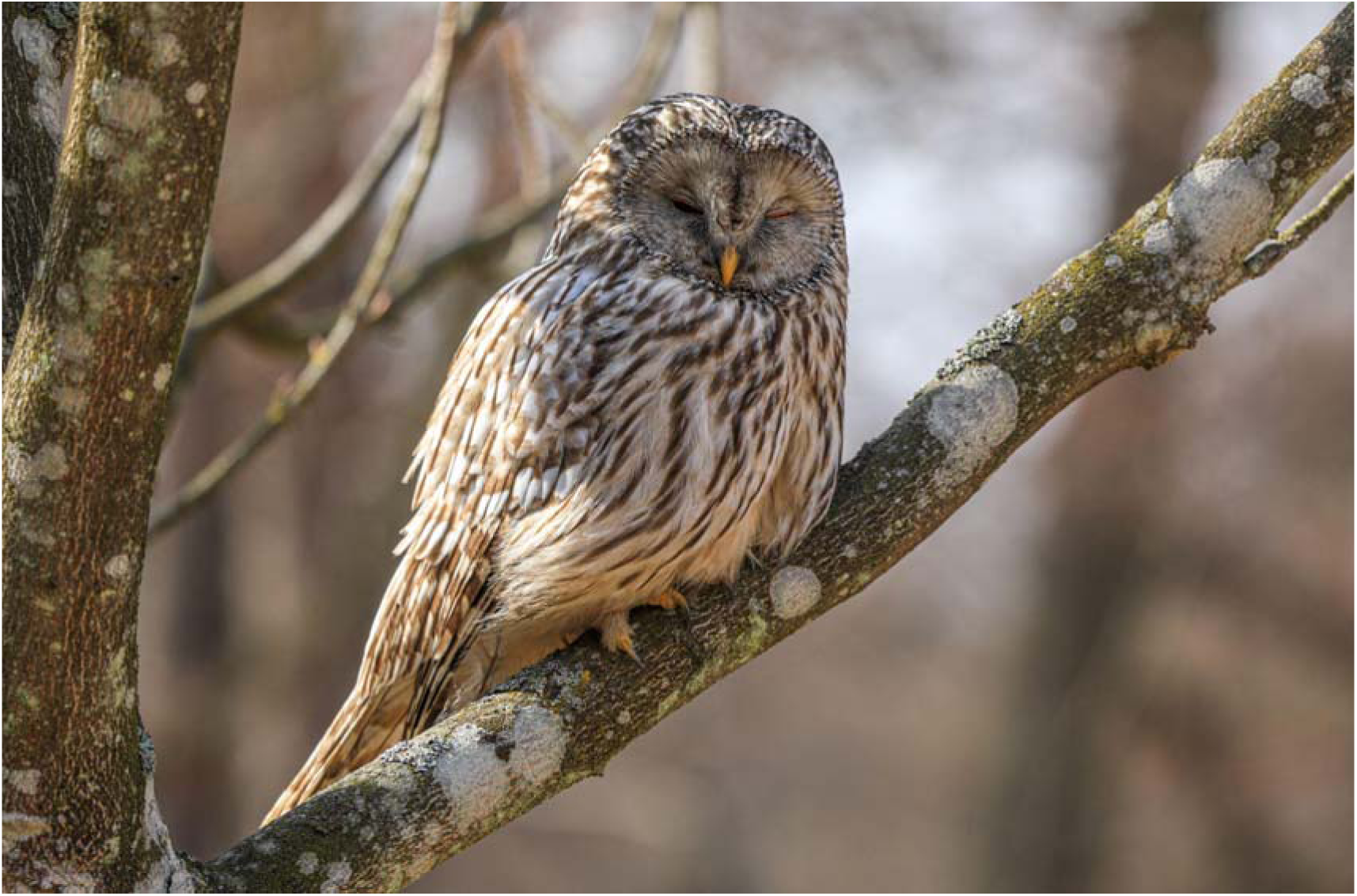
**A roosting Ural owl (*Strix uralensis*) of the reintroduced Central European population in the Viennese Forest. ©René Meißner**

The Ural owl is safeguarded under the EU Birds Directive Annex I, the Bern Convention on the Conservation of European Wildlife and Nature Habitats Annex I and II and listed on CITES Appendix II (see https://eunis.eea.europa.eu/species/1289). Although it is currently not considered a threatened species globally by the IUCN (BirdLife International, 2021), it has lost significant portions of its former range, particularly at its southwestern distribution edges. Historically, the species was also present in the low mountain ranges of Germany and the northern Alpine slopes of Switzerland, indicating a previously more continuous distribution that connected the Carpathian and Dinaric populations with the Northern European populations (Goffette et al., 2016). While still common in Fennoscandia, the loss of nesting sites due to forest management has resulted in threatened populations and local extinction in Southern Scandinavia and Central Europe (Scherzinger, 1987). Therefore, establishing new populations in the Ural owl’s former range and reinforcing existing ones is vital to ensure genetic exchange within the remaining metapopulations to secure the species ’long-term survival (Hausknecht et al., 2014). A crucial component of this effort is human-aided reintroduction, which has been conducted in Central Europe since the 1980s, for instance, in Germany and Austria (Engleder, 2003). Such projects aim to close distribution gaps and enable the Ural owl to recolonize suitable habitats on its own (Scherzinger, 1987).

To support these efforts, high-quality chromosome-level genome assemblies play an important role in species conservation (Paez et al., 2022) and provide an invaluable resource for informing and planning reintroduction programs (He et al., 2016). As highly contiguous genome references, assemblies heighten the value of short-read data, the most common genomic resource in conservation (B. R. Wright et al., 2020). Genomes further enable the assessment of genetic diversity, population structure, and subspecies, which are critical for maintaining the genetic health and adaptability of newly established populations (Formenti et al., 2022). Detecting inbreeding and emerging bottlenecks is especially important for reintroduction programs, as they guide the selection of individuals that maximize genetic variability (He et al., 2016). Furthermore, the detailed genetic information aids in understanding the species’ evolutionary history and adaptive traits, supporting the development of effective management strategies (Prost et al., 2022). Although fragmented scaffold assemblies might initially seem sufficient to provide genomic information comparable to chromosome-level assemblies for conservation genomic analyses, recent studies have shown that more contiguous assemblies enable better estimation of genetic diversity, improved localization, and visualization of regions with low heterozygosity within genomes, so-called runs of homozygosity, and allow the estimation of other important genetic parameters such as the amount of potentially harmful mutations in a genome, called mutational load (Totikov et al., 2021; von Seth et al., 2021). Here, we present a chromosome-level genome assembly, aiding our understanding of Ural owl populations and ultimately supporting conservation strategies for the species.

## Methods

### Biological Materials

In this study, we sequenced the genome of a Ural owl *Strix uralensis* (Voucher No. OV.35916 (http://id.zmuo.oulu.fi/OV.35916), GBIF entry: https://www.gbif.org/occurrence/4463254306) from muscle tissue stored at - 20°C at the frozen collection of the Zoological Museum of the University of Oulu. The female individual was found dead (famished) on the 29th of March 2001 in the municipality of Kuhmo near Lentiira, Finland, on the eastern side of southern Lake Änättijärvi.

In addition, we used tissue samples from two additional specimens from the frozen collection of the pathology department at the University of Veterinary Medicine Vienna, Austria, for RNA extraction: brain, kidney, muscle, and heart tissue from the female juvenile specimen AC/963/14 (3442), and liver tissue from the adult female specimen Z/677/21 (9679).

### Nucleic acid library preparation

We extracted high molecular weight (HMW) genomic DNA from the frozen muscle tissue of the sample (Voucher No. OV.35916) with the Monarch® HMW DNA Extraction Kit for Tissue (New England Biolabs, Ipswich, MA, USA) following the manufacturer’s protocol. The tissue was homogenized using the included Monarch Pestle Tubes and pestle. A second extraction was performed using the MagAttract HMW DNA kit (Qiagen, Hilden, Germany). The purity, quantity, and molecular weight of the DNA extract were checked using a NanoDrop Microvolume Spectrophotometer (Thermo Fisher Scientific Inc., Waltham, MA, USA), a Qubit Fluorometer with the Qubit Broad Range, dsDNA Quantification Assay Kit (Thermo Fisher Scientific Inc., Waltham, MA, USA), and an agarose gel electrophoresis, respectively.

In total, three long-read DNA libraries were prepared using the Oxford Nanopore Technologies (ONT, Oxford, UK) Ligation Sequencing Kit V14 (SQK-LSK114) with some modifications. Prior to library preparation, we sheared 2.9 µg of HMW DNA in a volume of 100 µl elution buffer by passing the solution through a 30G needle seven times, interspersed with a quick vortexing. We increased the incubation time during end-prep to 30 min for both temperatures and increased the elution times of the magnetic bead clean-up to 10 min after end-prep and 15 min at 37 °C after adapter ligation.

In addition, we generated a 150 bp paired-end chromatin conformation capture library from the frozen tissue using the Dovetail® Omni-C® Kit (Dovetail Genomics, part of Cantana Bio, LLC, Scotts Valley, CA, USA) and the NEBNext® Ultra™ II DNA Library Prep Kit for Illumina® according to the manufacturer’s protocols.

To generate mRNA evidence for the annotation of the genome assembly and to generate a transcriptome assembly, we extracted RNA from the brain, kidney, muscle, heart, and liver tissue with the Quick RNA Miniprep Plus Kit (Zymo Research, Orange, CA, USA). The extracted RNA was sent to Novogene Europe (Cambridge, UK) for 150 bp paired-end library preparation and sequencing.

### DNA/RNA Sequencing and Genome Assembly

From the three long-read ONT libraries, we initially sequenced two on the MinION Mk1c on a FLO-MIN114 R10.4.1 flow cell. The final library was sequenced on the PromethION 2 Solo (P2S) on a FLO-PRO114M R10.4.1 flow cell. The Omni-C library and the RNAseq libraries were sequenced at Novogene Europe (Cambridge, UK) on the Illumina Novaseq X platform. The raw ONT data were basecalled with Dorado v.0.5.2 (Oxford Nanopore Technologies Ltd., 2022) using the super-high-accuracy basecalling model in duplex mode. Read quality and length were checked with Nanoplot v.1.42 (De Coster et al., 2018).

The genome was assembled using the full ONT dataset (duplex + simplex reads) with Flye v.2.9.3 (Kolmogorov et al., 2019), including one iteration of long-read polishing. The contigs of the draft assembly were anchored into chromosome-scale scaffolds using the chromatin conformation information of the Omni-C data. We followed the Arima Hi-C mapping pipeline used by the Vertebrate Genome Project (https://github.com/VGP/vgp-assembly/blob/master/pipeline/salsa/arima_mapping_pipeline.sh) to filter and map the reads to the assembly. The reads were trimmed using fastp v.0.20.0 (Chen et al., 2018) and mapped to the assembly using bwa-mem v.0.7.17 (Li, 2013). Mapped reads were filtered based on mapping quality, read quality, and CIGAR strings with samtools v.1.18 (Li et al., 2009). Duplicated reads were removed using Sambamba v.1.0.1 (Tarasov et al., 2015). The mapped and filtered reads were subsequently used for proximity-ligation-based scaffolding with YaHS v1.1 (Zhou et al., 2022). Hi-C contact maps and assembly files used for manual curation in JuiceBox v.1.11.08 (Durand et al., 2016) were generated using JuicerTools v.1.22.01 (Durand et al., 2016). To improve the contig-level contiguity of the scaffolded assembly, we ran TGS-GapCloser v.2.0.0 (Xu et al., 2020) with the long-read ONT data followed by one iteration of short-read polishing with pilon v.1.24 (Walker et al., 2014) using the high-quality Omni-C data to improve base-level accuracy. We repeated scaffolding, gap-closing, and short-read polishing once to improve the assembly after manual corrections. In addition, we used GetOrganelle v.1.7.7.1 (Jin et al., 2020) to assemble the mitochondrial genome from the short-read Omni-C data.

#### Assembly QC & Synteny analyses

To assess the quality of the assembly, we calculated assembly statistics with Quast v.5.0.2 (Gurevich et al., 2013) and ran a gene set completeness analysis with both BUSCO v.5.4.7 (Manni et al., 2021) and compleasm v.0.2.6 (Huang & Li, 2023) using the aves_odb10 dataset. In addition, we estimated assembly completeness and the base-level error rate based on 21-mer counts generated with meryl v.1.4.1 using merqury v.1.3 (Rhie et al., 2020). To assess potential contamination, we first mapped the ONT long-reads, as well as the Omni-C illumina short-reads (separately) to the assembly using minimap2 v.2.28 (Li, 2018) and bwa-mem v.0.7.17 (Li, 2013), respectively and calculated mapping statistics with QualiMap v. 2.3 (Okonechnikov et al., 2016). We also used BLASTN v.2.11+ (Camacho et al., 2009) to assign taxon information to each of the scaffolds and contigs of the assembly. The resulting mapping files, as well as the output of BLASTN, were then combined into a blobplot with Blobtoolkit v. 3.5.2 (Challis et al., 2020). Synteny between the final Ural owl assembly and the available genome of *Strix occidentalis* (GCA_030819815.1), was analyzed with JupiterPlot v.1.1 (Chu, 2018).

#### Transcriptome assembly

The transcriptome of the Ural owl was assembled from the RNAseq data derived from the five different tissue samples. We first combined all RNAseq data into a single dataset before k-mer based read correction with Rcorrector v.1.0.7 (Song & Florea, 2015) and subsequent removal of all unfixable read pairs with the python script FilterUncorrectabledPEfastq.py (https://github.com/harvardinformatics/TranscriptomeAssemblyTools/). Next, we trimmed adaptors and low-quality bases from the filtered dataset using TrimGalore v.0.6.10 (Krueger, 2015) and removed unwanted rRNA reads (e.g., those from microorganisms) by mapping the data to the SILVA rRNA database v. 138.1 (Quast et al., 2013) using bowtie2 v.2.5.3 (Langmead & Salzberg, 2012), keeping only paired and unmapped reads.

This final dataset was then used to assemble the transcriptome using Trinity v.2.15 (Grabherr et al., 2011). To evaluate the quality of the transcriptome, we generated assembly statistics with the TrinityStats.pl script as part of Trinity and with Quast v.5.0.2 (Gurevich et al., 2013). Furthermore, we quantified read support by mapping the filtered reads back to the final assembly using bowtie2 and checked for completeness using BUSCO v.5.4.7 (Manni et al., 2021) in transcriptome mode and compleasm v.0.2.6 (Huang & Li, 2023).

#### Repeat and Gene annotation

Repeats in the genome assembly were masked in a three-step process. First, we masked known repeats for birds (‘*-species aves*’) based on the Repbase (release 20181026) (Bao et al., 2015) and Dfam (release 3.1-rb20181026) (Storer et al., 2021) databases with RepeatMasker v.4.1.0 (Smit et al., 2015a). Next, we identified the remaining repeats in the assembly *de novo* using RepeatModeler v.2.0.1 (Smit et al., 2015b). The resulting *de novo* repeat library was used in a second iteration of repeat masking to mask the remaining repeats in the assembly. We combined both Repeatmasker repeat tables into a single table to represent the entirety of masked repeats in the assembly (Supplementary Material S1).

Genes in the masked assembly were predicted based on homology with

GeMoMa v.1.9 (Keilwagen et al., 2018) using the following eight annotated assemblies as evidence: Chicken (*Gallus gallus*) GCF_016699485.2, Japanese quail (*Coturnix japonica*) GCF_001577835.2, Burrowing owl (*Athene cunicularia*) GCF_003259725.1 (Mueller et al., 2018), Common barn owl (*Tyto alba*) GCF_018691265.1 (Cumer et al., 2022), Speckled mousebird (*Colius striatus*) GCF_028858725.1, California Condor (*Gymnogypus californianus*) GCF_018139145.2 (Robinson et al., 2021), Red-fronted tinkerbird (*Pogoniulus pusillus*) GCF_015220805.1, Downy woodpecker (*Dryobates pubescens*) GCF_014839835.1. In addition, the corrected and trimmed RNAseq data of the five different tissues were mapped against the masked reference with STAR v.2.7.9a (Dobin et al., 2013) and used as extrinsic evidence during the annotation. The proteins predicted by GeMoMa were further annotated by a BLASTP v2.15.0+ (Camacho et al., 2009) search against the SwissProt database (release 04-2024, The UniProt Consortium, 2019) applying a *e*-value cutoff of 10^-6^. We also annotated GeneOntology (GO) terms, domains, and motifs with InterProScan v.5.64-96.0 (Jones et al., 2014).

## Results

### Genome sequencing and assembly

Sequencing on the ONT MinION Mk1c and PromethION P2S together generated a total of 49.24 Gb or approximately 39-fold sequencing depth of long-read data after base-calling, of which 37.37 Gb were simplex, and 5.74 Gb were duplex reads. The combined long-read dataset had a mean base quality of 14.7, a median base quality of 20.1, and a mean read length of 7,209.9 bp.

The final assembly (*S3B_Suralensis_v1.3.3*) after initial assembly with Flye, proximity-ligation scaffolding, gap-closing, and removal of potential contamination, had a total length of 1.26 Gb across 1,876 scaffolds/contigs, including one contig for the mitochondrial genome, with scaffold and contig N50 values of 88.65 Mb and 19.98 Mb, respectively, and no remaining signs of contamination (Table 2A, Fig. 2 A-C). The largest 42 scaffolds, likely corresponding to the expected number of haploid chromosomes (including a partial W chromosome) (Sasaki et al., 1994), contain 96.42 % of the total assembly length, resulting in an L50 of five. *S3B_Suralensis_v1.3.3* showed high gene set completeness with BUSCO and compleasm scores of 97.5 % and 99.65 % complete BUSCO genes of the aves_odb10 dataset with only 0.5% and 0.28 % duplicated genes, respectively (Table 2B). Furthermore, Merqury estimated a k-mer completeness of 95.18 % with a QV score of 39.45, corresponding to an error rate of 0.0001.

**Figure 2.**
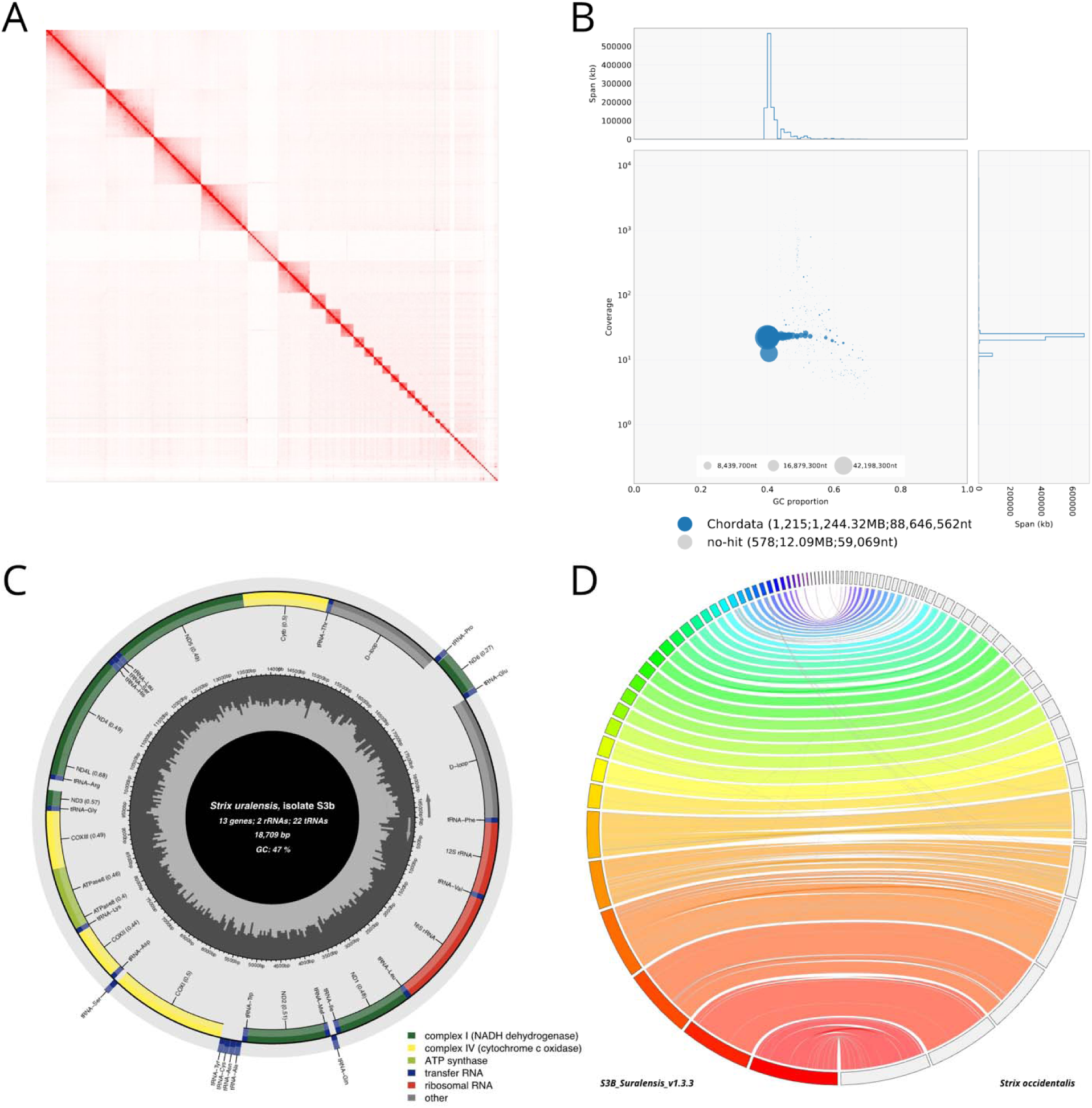
**Assembly quality assessment and mitochondrial genome of *S3B_Suralensis_v1.3.3* and synteny with *S. occidentalis*.** (A)Hi-C (Omni-C) contact density map depicting the 42 chromosome-level scaffolds and additional small scaffolds of *S3B_Suralensis_v1.3.3*. (B) BlobPlot analysis comparing GC content (x-axis) and sequence coverage of the Omni-C data (y-axis). The color of the “blobs” representing each scaffold corresponds to the taxonomic assignment based on NCBI’s nucleotide database. (C) Graphical representation of the annotated mitochondrial genome of *S. uralensis* isolate S3B. (D) Circos plot generated with JupiterPlot comparing the synteny of *S3B_Suralensis_v1.3.3* (left) with its close relative *S. occidentalis* (right). Colored ribbons between scaffolds indicate syntenic regions. Scaffolds are sorted by size from the largest (bottom) to the smallest (top).

**Table 2.** Assembly statistics and gene set completeness scores.

**A.** Assembly statistics of *S3B\_Suralensis\_v1.3.3* and the *Strix uralensis* transcriptome
|  | Scaffold-level | Contig-level* | Transcriptome |
| --- | --- | --- | --- |
|  | *S3B_Suralensis_v1.3.3* | *S3B_Suralensis_v1.3.3\** |  |
| **No. of scaffolds/contigs** | 1,793 | 1,985 | 316,288 |
| **No. of scaffolds/contigs (> 1 kbp)** | 1,611 | 1,811 | 74,767 |
| **L50** | 5 | 10 | 23,310 |
| **N50 (bp)** | 88,646,562 | 21,740,082 | 3,587 |
| **Max. scaffold/contig length (bp)** | 168,792,956 | 164,885,455 | 29,414 |
| **Total length (bp)** | 1,256,413,995 | 1,256,394,007 | 332,292,401 |
| **GC (%)** | 42.27 | 42.27 | 46.45 |
| **No. of N's** | 19,988 | 0 | 221 |
| **No. of N's per 100 kbp** | 1.59 | 0 | 0.08 |

\*broken into contigs at gaps with a length of $\geq 10$ N's. Statistics for these columns are based on contigs, the remaining columns are based on scaffolds.

B. Gene set completeness scores derived from BUSCO and compleasm
| **Assembly** | **Tool** | **Single copy (%)** | **Duplicated (%)** | **Fragmented (%)** | **Missing (%)** |
| --- | --- | --- | --- | --- | --- |
| **S3B_Suralensis_v1.3.3** | **BUSCO** | 97.0 (8085) | 0.5 (45) | 0.4 (33) | 2.1 (175) |
|  | **compleasm** | 99.65 (8308) | 0.28 (23) | 0.00 (0) | 0.07 (6) |
| **S3B_Suralensis_v1.3.3**<br>**largest 42 scaffolds** | **BUSCO** | 97.0 (8085) | 0.5 (38) | 0.4 (35) | 2.1 (180) |
|  | **compleasm** | 99.62 (8305) | 0.25 (21) | 0.00 (0)* | 0.13 (11) |
| **Annotation** | **BUSCO** | 44.0 (3671) | 53.3 (4448) | 0.7 (59) | 2.0 (160) |
|  | **compleasm** | 38.68 (3225) | 58.60 (4886) | 0.82 (68) | 1.91 (159) |
| **Transcriptome** | **BUSCO** | 28.0 (2337) | 60.2 (5021) | 2.8 (235) | 9.0 (745) |
|  | **compleasm** | 33.07 (2757) | 55.09 (4593) | 3.30 (275)* | 8.54 (712) |

\*This score includes both "Fragmented Genes" from subclass 1 and subclass 2

### Transcriptome assembly

The final transcriptome assembly based on 39.7 Gb of RNAseq data has a total length of 333.2 Mb with a contig N50 of 2,792 bp, a total number of trinity ‘genes ’ and transcripts of 241,711 and 318,498, respectively (Table 2A). BUSCO found 88.2 % complete orthologous genes of the aves_odb10 dataset, with 28.0 % being single copy, 60.2 % duplicated, and 9.0 % missing (Table 2B). Compleasm identified 88.16% complete genes, of which 33.07 % were single-copy, 55,09 % duplicate genes, and 8.54% missing genes (Table 2B).

### Annotation Repeat annotation

A total of 156.83 Mb (12.48 %) of the final assembly was classified as repeats (Supplementary Material S1). Specifically, interspersed repeats make up most of the repeats (10.39 %), of which Long Interspersed Nuclear Elements (LINEs) are the most common repeat elements at 4,90 %, followed by Long Terminal Repeats (LTRs) with 1.92 %. An additional 2.91 % of the assembly was identified as unknown or unclassified interspersed repeats. Of the non-interspersed repeats, simple repeats are the most common, spanning 1.14 % of the assembly.

### Gene annotation

The homology-based gene prediction with GeMoMa identified 17,650 genes spanning 328.2 Mb of the assembly, with a median gene length of 8,985 bp. BUSCO and compleasm analyses indicate high completeness of the annotation with 97.3 % complete orthologous (single copy and duplicates) of the aves_odb10 dataset found in the annotation with BUSCO and 96.77 % identified by compleasm (Table 2B). Of the 44,008 predicted proteins, 43,016 (97.74 %) could be matched to entries within the Swiss-Prot database, while InterProScan assigned a functional annotation to 43,842 (99.62%) proteins. At least one Gene Ontology (GO)-term was assigned to 33,608 (76.37 %) proteins, and 39,328 (89.37 %) proteins were assigned to the reactome.

## Discussion

The high-quality near chromosome-level assembly of the Ural owl presented in this study represents a valuable genomic resource for the species and the Stringiformes. With highly contiguous scaffolds and gene set completeness, the assembly offers a crucial genomic base for future population and conservation genetics studies (B. Wright et al., 2019) and could help to shed light on the still difficult to resolve phylogenetic position of the Strigiformes within the Telluraves (Stiller et al., 2024). The assembly’s quality is evidenced by its high BUSCO scores and robust annotation, which highlights its suitability for in-depth genetic analysis and its role in understanding complex evolutionary processes. Moreover, detailed gene annotation enables future studies on gene function and the species ’adaptive capacity (King et al., 2003).

Furthermore, this genome will greatly aid Ural owl conservation and provide insights into genetic diversity, population structure, and adaptive traits, key factors in evaluating wild populations and identifying potential source populations for successful reintroduction programs (He et al., 2016). Similar to the reference genomes developed for other non-model species, this Ural owl assembly will support a range of further applications, including identifying genes linked to adaptive traits in the species, monitoring population-level genetic diversity, and ensuring the genetic health of reintroduced populations (He et al., 2016; Paez et al., 2022). High-quality references are essential for developing effective conservation strategies, especially for species experiencing habitat loss or fragmentation.

Although obtaining high-quality genome assemblies for non-model species can be challenging due to technical and financial limitations, advances in sequencing technologies and bioinformatics make these resources more accessible and economically feasible (McMahon et al., 2014). Collaboration between institutions and global genome initiatives can further accelerate the development of genomic resources for conservation purposes (Formenti et al., 2022). Such collaborations allow conservation managers to focus on adaptive management while generating high-quality genomic data to inform real-time decision-making (Bernos et al., 2020).

In summary, the new Ural owl genome assembly provides an essential tool for conservation efforts by enabling detailed genetic analysis and population monitoring. This resource will play a key role in supporting species reintroduction and long-term population viability, ensuring genetic diversity, and identifying adaptive traits critical for the species’ future survival in the wild. As global biodiversity faces increasing threats, the development and application of genomic resources will be crucial in shaping effective conservation strategies for endangered species and will aid in impeding global biodiversity loss.

## Acknowledgments

We thank Anna Kübber-Heiss and Helmut Dier from the pathology department of the University of Veterinary Medicine, Vienna, for access to their frozen tissue sample collection, and Pasi Laakso and Ilmari Mäkisalo for collecting and providing the reference specimen in Kuhmo, Finland.

## Data Availability

All primary data (DNA/RNA sequences) underlying these analyses, the genome assembly, and the transcriptome have been deposited under GenBank BioProject PRJNA1140424. The final genome assembly, transcriptome assembly, repeat masked assembly and annotation results are deposited at Dryad (*a link will be provided during the revision*).

---

**Table 1.** Software and versions used to generate the *Strix uralensis* assembly and transcriptome.

| *Pipeline Step* | *Software* | *Version* | *RRID* |
| --- | --- | --- | --- |
| ONT Basecalling | Dorado | v.0.5.2 | SCR 025883 |
| ONT QC | NanoPlot | v.1.42 | SCR 024128 |
| Short-read trimming and filtering | fastp | v.0.23.4 | SCR 016962 |
| Assembly & long-read polishing | Flye | v.2.9.3 | SCR 017016 |
| Mapping of short reads | BWA-MEM | v.0.7.17 | SCR 010910 |
| Mapping of long reads | minimap2 | v.2.28 | SCR 018550 |
| Sort & index BAM | samtools | v.1.18 | SCR 002105 |
| Duplicate removal | sambamba | v.1.0.1 | SCR 024328 |
| Hi-C data processing for scaffolding | Arima Genomics mapping pipeline | Commit 2e74ea4 |  |
| Scaffolding | YaHS | v.1.1 | SCR 022965 |
| Hi-C contact map generation | JuicerTools | v.1.22.01 | SCR 017226 |
| Manual editing of Hi-C contact map | Juicebox | v.1.11.08 | SCR 021172 |
| Gap-closing | TGS-GapCloser | v.2.0.0 | SCR 017633 |
| Short-read polishing | Pilon | v.1.24 | SCR 014731 |
| Assembly statistics | Quast | v.5.0.2 | SCR 001228 |
| Gene set completeness | BUSCO | v.5.4.7 | SCR 015008 |
| Gene set completeness | compleasm | v.0.2.6 | SCR 026370 |
| Mapping statistics | QualiMap | v.2.3 | SCR 001209 |
| Assembly completeness | Merqury | v.1.3 | SCR 022964 |
| Contamination analyses | BLASTN | v.2.11.0+ | SCR 001598 |
| Contamination analyses | BlobToolKit | v.3.5.2 | SCR 025882 |
| Mitochondrial genome assembly | GetOrganelle | v.1.7.7.1 | SCR 022963 |
| RNAseq read correction | Rcorrector | v.1.0.7 | SCR 022011 |
| RNAseq read adaptor trimming | Trim Galore | v.0.6.10 | SCR 011847 |
| RNAseq mapping to remove rRNA | Bowtie2 | v.2.5.3 | SCR_016368 |
| Transcriptome assembly | Trinity | v.2.15.2 | SCR_013048 |
| RNAseq data mapping to genome | STAR | v.2.7.9a | SCR_004463 |
| Repeat library | RepeatModeler | v.2.0.1 | SCR_015027 |
| Repeat masking | RepeatMasker | v.4.1.0 | SCR_012954 |
| Homology-based gene prediction | GeMoMa | v.1.9 | SCR_017646 |
| Functional annotation | InterProScan | v.5.64.90 | SCR_005829 |
| Functional annotation | BLASTP | v.2.15.0+ | SCR_001010 |
| Synteny analyses | JupiterPlot | v.1.1 | SCR_022961 |

## Supplementary Material

**S1:** Repeat table generated by RepeatMasker

## References

Ackerman, J. (2024). What an Owl Knows: The New Science of the World’s Most Enigmatic Birds. Penguin Group.

Bao, W., Kojima, K. K., & Kohany, O. (2015). Repbase Update, a database of repetitive elements in eukaryotic genomes. Mobile DNA, 6(1), 11. 10.1186/s13100-015-0041-9

Bernos, T. A., Jeffries, K. M., & Mandrak, N. E. (2020). Linking genomics and fish conservation decision making: A review. Reviews in Fish Biology and Fisheries, 30(4), 587–604. 10.1007/s11160-020-09618-8

BirdLife International. (2021). IUCN Red List of Threatened Species: Strix uralensis. *IUCN Red List of Threatened Species*. https://www.iucnredlist.org/en

Camacho, C., Coulouris, G., Avagyan, V., Ma, N., Papadopoulos, J., Bealer, K., & Madden, T. L. (2009). BLAST+: Architecture and applications. BMC Bioinformatics, 10(1), 421. 10.1186/1471-2105-10-421

Challis, R., Richards, E., Rajan, J., Cochrane, G., & Blaxter, M. (2020). BlobToolKit – Interactive Quality Assessment of Genome Assemblies. G3 Genes|Genomes|Genetics, 10(4), 1361–1374. 10.1534/g3.119.400908

Chen, S., Zhou, Y., Chen, Y., & Gu, J. (2018). fastp: An ultra-fast all-in-one FASTQ preprocessor. Bioinformatics, 34(17), i884–i890. 10.1093/bioinformatics/bty560

Chu, J. (2018). Jupiter Plot: A Circos-based tool to visualize genome assembly consistency. 10.5281/zenodo.1241235

Cumer, T., Machado, A. P., Siverio, F., Cherkaoui, S. I., Roque, I., Lourenço, R., Charter, M., Roulin, A., & Goudet, J. (2022). Genomic basis of insularity and ecological divergence in barn owls (Tyto alba) of the Canary Islands. Heredity, 129(5), 281–294. 10.1038/s41437-022-00562-w

De Coster, W., D’Hert, S., Schultz, D. T., Cruts, M., & Van Broeckhoven, C. (2018). NanoPack: Visualizing and processing long-read sequencing data. Bioinformatics, 34(15), 2666–2669. 10.1093/bioinformatics/bty149

Dobin, A., Davis, C. A., Schlesinger, F., Drenkow, J., Zaleski, C., Jha, S., Batut, P., Chaisson, M., & Gingeras, T. R. (2013). STAR: Ultrafast universal RNA-seq aligner. *Bioinformatics (Oxford*, England), 29(1), 15–21. 10.1093/bioinformatics/bts635

Durand, N. C., Shamim, M. S., Machol, I., Rao, S. S. P., Huntley, M. H., Lander, E. S., & Aiden, E. L. (2016). Juicer Provides a One-Click System for Analyzing Loop-Resolution Hi-C Experiments. Cell Systems, 3(1), 95–98. 10.1016/j.cels.2016.07.002

Engleder, T. (2003). Re-introduction of the Ural Owl (Strix uralensis) on the Austrian side of the Bohemian Forest in 2001. Buteo, 13, 97–99.

Formenti, G., Theissinger, K., Fernandes, C., Bista, I., Bombarely, A., Bleidorn, C., Ciofi, C., Crottini, A., Godoy, J. A., Höglund, J., Malukiewicz, J., Mouton, A., Oomen, R. A., Paez, S., Palsbøll, P. J., Pampoulie, C., Ruiz-López, M. J., Svardal, H., Theofanopoulou, C., … Zammit, G. (2022). The era of reference genomes in conservation genomics. Trends in Ecology & Evolution, 37(3), 197–202. 10.1016/j.tree.2021.11.008

Goffette, Q., Denis, M., Pöllath, N., & Neer, W. V. (2016). Change in Historical Range of the Ural Owl in Europe. Belgian Journal of Zoology, 146(1), Article 1. 10.26496/bjz.2016.37

Grabherr, M. G., Haas, B. J., Yassour, M., Levin, J. Z., Thompson, D. A., Amit, I., Adiconis, X., Fan, L., Raychowdhury, R., Zeng, Q., Chen, Z., Mauceli, E., Hacohen, N., Gnirke, A., Rhind, N., Palma, F. di, Birren, B. W., Nusbaum, C., Lindblad-Toh, K., … Regev, A. (2011). Full-length transcriptome assembly from RNA-Seq data without a reference genome. Nature Biotechnology, 29(7), 644–652. 10.1038/nbt.1883

Gurevich, A., Saveliev, V., Vyahhi, N., & Tesler, G. (2013). QUAST: Quality assessment tool for genome assemblies. Bioinformatics, 29(8), 1072–1075. 10.1093/bioinformatics/btt086

Hausknecht, R., Jacobs, S., Müller, J., Zink, R., Frey, H., Solheim, R., Vrezec, A., Kristin, A., Mihok, J., Kergalve, I., Saurola, P., & Kuehn, R. (2014). Phylogeographic analysis and genetic cluster recognition for the conservation of Ural Owls (Strix uralensis) in Europe. Journal of Ornithology, 155(1), 121–134. 10.1007/s10336-013-0994-8

He, X., Johansson, M. L., & Heath, D. D. (2016). Role of genomics and transcriptomics in selection of reintroduction source populations. Conservation Biology, 30(5), 1010–1018. 10.1111/cobi.12674

Huang, N., & Li, H. (2023). compleasm: A faster and more accurate reimplementation of BUSCO. Bioinformatics, 39(10), btad595. 10.1093/bioinformatics/btad595

Jin, J.-J., Yu, W.-B., Yang, J.-B., Song, Y., dePamphilis, C. W., Yi, T.-S., & Li, D.-Z. (2020). GetOrganelle: A fast and versatile toolkit for accurate de novo assembly of organelle genomes. Genome Biology, 21(1), 241. 10.1186/s13059-020-02154-5

Jones, P., Binns, D., Chang, H.-Y., Fraser, M., Li, W., McAnulla, C., McWilliam, H., Maslen, J., Mitchell, A., Nuka, G., Pesseat, S., Quinn, A. F., Sangrador-Vegas, A., Scheremetjew, M., Yong, S.-Y., Lopez, R., & Hunter, S. (2014). InterProScan 5: Genome-scale protein function classification. Bioinformatics, 30(9), 1236–1240. 10.1093/bioinformatics/btu031

Keilwagen, J., Hartung, F., Paulini, M., Twardziok, S. O., & Grau, J. (2018). Combining RNA-seq data and homology-based gene prediction for plants, animals and fungi. BMC Bioinformatics, 19(1). 10.1186/s12859-018-2203-5

King, O. D., Foulger, R. E., Dwight, S. S., White, J. V., & Roth, F. P. (2003). Predicting Gene Function From Patterns of Annotation. Genome Research, 13(5), 896–904. 10.1101/gr.440803

Kolmogorov, M., Yuan, J., Lin, Y., & Pevzner, P. A. (2019). Assembly of long, error-prone reads using repeat graphs. Nature Biotechnology, 37(5), 540–546. 10.1038/s41587-019-0072-8

Konishi, M. (1973). How the Owl Tracks Its Prey: Experiments with trained barn owls reveal how their acute sense of hearing enables them to catch prey in the dark. American Scientist, 61(4), 414–424.

Korpimäki, E., & Sulkava, S. (1987). Diet and breeding performance of Ural Owls Strix uralensis under fluctuating food conditions. Ornis Fennica, 64(2), 57– 66.

Krueger, F. (2015). Trim Galore!: A wrapper around Cutadapt and FastQC to consistently apply adapter and quality trimming to FastQ files, with extra functionality for RRBS data. *Babraham Institute*.

Langmead, B., & Salzberg, S. L. (2012). Fast gapped-read alignment with Bowtie 2. Nature Methods, 9(4), 357–359. 10.1038/nmeth.1923

Li, H. (2013). Aligning sequence reads, clone sequences and assembly contigs with BWA-MEM. arXiv:1303.3997 [q-Bio]. 10.48550/arXiv.1303.3997

Li, H. (2018). Minimap2: Pairwise alignment for nucleotide sequences. Bioinformatics, 34(18), 3094–3100. 10.1093/bioinformatics/bty191

Li, H., Handsaker, B., Wysoker, A., Fennell, T., Ruan, J., Homer, N., Marth, G., Abecasis, G., & Durbin, R. (2009). The Sequence Alignment/Map format and SAMtools. Bioinformatics, 25(16), 2078–2079. 10.1093/bioinformatics/btp352

Manni, M., Berkeley, M. R., Seppey, M., Simão, F. A., & Zdobnov, E. M. (2021). BUSCO Update: Novel and Streamlined Workflows along with Broader and Deeper Phylogenetic Coverage for Scoring of Eukaryotic, Prokaryotic, and Viral Genomes. Molecular Biology and Evolution, 38(10), 4647–4654. 10.1093/molbev/msab199

McMahon, B. J., Teeling, E. C., & Höglund, J. (2014). How and why should we implement genomics into conservation? Evolutionary Applications, 7(9), 999–1007. 10.1111/eva.12193

Mueller, J. C., Kuhl, H., Boerno, S., Tella, J. L., Carrete, M., & Kempenaers, B. (2018). Evolution of genomic variation in the burrowing owl in response to recent colonization of urban areas. *Proceedings*. Biological Sciences, 285(1878), 20180206. 10.1098/rspb.2018.0206

Okonechnikov, K., Conesa, A., & García-Alcalde, F. (2016). Qualimap 2: Advanced multi-sample quality control for high-throughput sequencing data. Bioinformatics, 32(2), 292–294. 10.1093/bioinformatics/btv566

Oxford Nanopore Technologies Ltd. (2022). *Dorado* [Computer software]. Oxford Nanopore Technologies. https://github.com/nanoporetech/dorado

Paez, S., Kraus, R. H. S., Shapiro, B., Gilbert, M. T. P., Jarvis, E. D., & VERTEBRATE GENOMES PROJECT CONSERVATION GROUP. (2022). Reference genomes for conservation. Science, 377(6604), 364–366. 10.1126/science.abm8127

Penhallurick, J. M. (2002). The taxonomy and conservation status of the owls of the world: A review. *Ecology and Conservation of Owls. CSIRO Publishing*, Collingwood, 343–354.

Prost, S., Machado, A. P., Zumbroich, J., Preier, L., Mahtani-Williams, S., Meissner, R., Guschanski, K., Brealey, J. C., Fernandes, C. R., Vercammen, P., Hunter, L. T. B., Abramov, A. V., Plasil, M., Horin, P., Godsall-Bottriell, L., Bottriell, P., Dalton, D. L., Kotze, A., & Burger, P. A. (2022). Genomic analyses show extremely perilous conservation status of African and Asiatic cheetahs (Acinonyx jubatus). Molecular Ecology, 31(16), 4208–4223. 10.1111/mec.16577

Quast, C., Pruesse, E., Yilmaz, P., Gerken, J., Schweer, T., Yarza, P., Peplies, J., & Glöckner, F. O. (2013). The SILVA ribosomal RNA gene database project: Improved data processing and web-based tools. Nucleic Acids Research, 41(D1), D590–D596. 10.1093/nar/gks1219

Rhie, A., Walenz, B. P., Koren, S., & Phillippy, A. M. (2020). Merqury: Reference-free quality, completeness, and phasing assessment for genome assemblies. Genome Biology, 21(1), 245. 10.1186/s13059-020-02134-9

Robinson, J. A., Bowie, R. C. K., Dudchenko, O., Aiden, E. L., Hendrickson, S. L., Steiner, C. C., Ryder, O. A., Mindell, D. P., & Wall, J. D. (2021). Genome-wide diversity in the California condor tracks its prehistoric abundance and decline. Current Biology: CB, 31(13), 2939–2946.e5. 10.1016/j.cub.2021.04.035

Sasaki, M., Nishida-Umehara, C., & Tsuchiya, K. (1994). A Comparative Study of G-banded Karyotypes in Eight Species of Owls. Cytologia, 59(2), 183–185. 10.1508/cytologia.59.183

Scherzinger, W. T. (1987). Reintroduction of the ural owl in the Bavarian National Park, Germany. Biology and Conservation of Northern Forest Owls. USDA Forest Service GTR RM-142. Ft. Collins, CO USA, 75–80.

Smit, A., Hubley, R., & Green, P. (2015a). RepeatMasker Open-4.0. 2013–2015. Smit, A., Hubley, R., & Green, P. (2015b). RepeatModeler Open-1.0. 2008–2015. Seattle, USA: *Institute for Systems Biology*.

Song, L., & Florea, L. (2015). Rcorrector: Efficient and accurate error correction for Illumina RNA-seq reads. GigaScience, 4(1), s13742–015-0089-y. 10.1186/s13742-015-0089-y

Stiller, J., Feng, S., Chowdhury, A.-A., Rivas-González, I., Duchêne, D. A., Fang, Q., Deng, Y., Kozlov, A., Stamatakis, A., Claramunt, S., Nguyen, J. M. T., Ho, S. Y. W., Faircloth, B. C., Haag, J., Houde, P., Cracraft, J., Balaban, M., Mai, U., Chen, G., … Zhang, G. (2024). Complexity of avian evolution revealed by family-level genomes. Nature, 629(8013), 851–860. 10.1038/s41586-024-07323-1

Storer, J., Hubley, R., Rosen, J., Wheeler, T. J., & Smit, A. F. (2021). The Dfam community resource of transposable element families, sequence models, and genome annotations. Mobile DNA, 12(1), 2. 10.1186/s13100-020-00230-y

Tarasov, A., Vilella, A. J., Cuppen, E., Nijman, I. J., & Prins, P. (2015). Sambamba: Fast processing of NGS alignment formats. Bioinformatics, 31(12), 2032–2034. 10.1093/bioinformatics/btv098

The UniProt Consortium. (2019). UniProt: A worldwide hub of protein knowledge. Nucleic Acids Research, 47(D1), D506–D515. 10.1093/nar/gky1049

Totikov, A., Tomarovsky, A., Prokopov, D., Yakupova, A., Bulyonkova, T., Derezanin, L., Rasskazov, D., Wolfsberger, W. W., Koepfli, K.-P., Oleksyk, T. K., & Kliver, S. (2021). Chromosome-Level Genome Assemblies Expand Capabilities of Genomics for Conservation Biology. Genes, 12(9), Article 9. 10.3390/genes12091336

Tutiš, V., Radović, D., Ćiković, D., Barišić, S., & Kralj, J. (2009). Distribution, Density and Habitat Relationships of the Ural Owl Strix uralensis macroura in Croatia. Ardea, 97(4), 563–570. 10.5253/078.097.0423

von Seth, J., Dussex, N., Díez-del-Molino, D., van der Valk, T., Kutschera, V. E., Kierczak, M., Steiner, C. C., Liu, S., Gilbert, M. T. P., Sinding, M.-H. S., Prost, S., Guschanski, K., Nathan, S. K. S. S., Brace, S., Chan, Y. L., Wheat, C. W., Skoglund, P., Ryder, O. A., Goossens, B., … Dalén, L. (2021). Genomic insights into the conservation status of the world’s last remaining Sumatran rhinoceros populations. Nature Communications, 12(1), 2393. 10.1038/s41467-021-22386-8

Voous, K. H. (1964). Wood owls of the genera Strix and Ciccaba. Zoologische Mededelingen, 39(46), 471–478.

Walker, B. J., Abeel, T., Shea, T., Priest, M., Abouelliel, A., Sakthikumar, S., Cuomo, C. A., Zeng, Q., Wortman, J., Young, S. K., & Earl, A. M. (2014). Pilon: An Integrated Tool for Comprehensive Microbial Variant Detection and Genome Assembly Improvement. PLOS ONE, 9(11), e112963. 10.1371/journal.pone.0112963

Wright, B., Farquharson, K. A., McLennan, E. A., Belov, K., Hogg, C. J., & Grueber, C. E. (2019). From reference genomes to population genomics: Comparing three reference-aligned reduced-representation sequencing pipelines in two wildlife species. BMC Genomics, 20(1), 453. 10.1186/s12864-019-5806-y

Wright, B. R., Farquharson, K. A., McLennan, E. A., Belov, K., Hogg, C. J., & Grueber, C. E. (2020). A demonstration of conservation genomics for threatened species management. Molecular Ecology Resources, 20(6), 1526– 1541. 10.1111/1755-0998.13211

Xu, M., Guo, L., Gu, S., Wang, O., Zhang, R., Peters, B. A., Fan, G., Liu, X., Xu, X., Deng, L., & Zhang, Y. (2020). TGS-GapCloser: A fast and accurate gap closer for large genomes with low coverage of error-prone long reads. GigaScience, 9(giaa094). 10.1093/gigascience/giaa094

Zhou, C., McCarthy, S. A., & Durbin, R. (2022). YaHS: Yet another Hi-C scaffolding tool. Bioinformatics, 39(1), btac808. 10.1093/bioinformatics/btac808

